# *Drosophila melanogaster* infected with *Wolbachia* strain *w*MelCS prefer cooler temperatures

**DOI:** 10.1101/352872

**Authors:** Pieter A. Arnold, Samantha C. Levin, Aleksej L. Stevanovic, Karyn N. Johnson

## Abstract

1. Temperature plays a fundamental role in the dynamics of host-pathogen interactions.
2. *Wolbachia* is an endosymbiotic bacteria that infects about 40% of arthropod species, which can affect host behaviour and reproduction. Yet, the effect of *Wolbachia* on host thermoregulatory behaviour is largely unknown, despite its use in disease vector control programs in thermally variable environments.
3. Here, we used a thermal gradient to test whether *Drosophila melanogaster* infected with *Wolbachia* strain *w*MelCS exhibit different temperature preferences (*T*_p_) to uninfected flies.
4. We found that *Wolbachia*-infected flies preferred a cooler mean temperature (*T*_p_ = 25.06±0.25°C) than uninfected flies (*T*_p_ = 25.78±0.24°C).
5. This finding suggests that *Wolbachia*-infected hosts might seek out cooler microclimates to reduce exposure to and lessen the consequences of high temperatures. This finding has generated hypotheses that will be fruitful in areas of research for exploring the mechanisms by which the change in *T*_p_ occurs in this complex and significant host-pathogen-environment interaction.

## Introduction

*Wolbachia pipientis* is an endosymbiont bacteria that infects an estimated 40% of terrestrial arthropod species (Zug & Hammerstein, 2012). The association between *Wolbachia* and its hosts has been the subject of a wide array of studies, including the alteration of host behaviours and reproduction (Weeks et al., 2002), cytoplasmic incompatibility for disease vector control (Clancy & Hoffmann, 1998), and environmental factors mediating host-pathogen interactions (Murdock et al., 2012). Temperature is a key environmental modulator of host-pathogen interactions, which constrains the rate of biological reactions and sets limits to performance and survival (Thomas & Blanford, 2003).

For the insect host, there is little physiological capacity to differentiate their body temperature (*T*_b_) from the ambient temperature of their surrounding environment (Angilletta, 2009). Physiological rates and performance are strongly affected by *T*_b_ in ectotherms, so organisms should aim to maintain their *T*_b_ across a range of temperatures that correspond to adequate performance (Huey & Kingsolver, 1989). One strategy that ectotherms can employ to avoid exposure to unsuitable temperatures is to modify their behaviour to seek more suitable microclimates, such as in shade, to find their preferred temperature (*T*_p_) (Sunday et al., 2014). Temperature preference is the perception and neural integration of thermal information, resulting in this crucial thermoregulatory behaviour (Abram et al., 2017). *Drosophila melanogaster* exhibits strong circadian and neurally controlled temperature preference behaviour, which centres around 24–27°C (Arnold et al., 2015; Kaneko et al., 2012).

The thermal biology of *Wolbachia* and host associations is complex and variable. High temperatures are unfavourable for some strains, where *Wolbachia* density is much higher at lower temperatures (e.g., 13–19°C (Moghadam et al., 2017)) and is reduced at higher temperatures (e.g., 26°C (Clancy & Hoffmann, 1998)). *Wolbachia* can also be mostly or completely eliminated by exposure to cyclic heat stress or temperatures above 30°C, reducing vertical transmission of the symbiont (Corbin et al., 2016; Ross et al., 2017). However, higher temperatures do not always reduce *Wolbachia* titre and responses to temperature vary greatly among study systems (e.g., Mouton et al., 2006; Murdock et al., 2014). The *Wolbachia* strain *w*MelCS that we use here infects natural *D. melanogaster* populations, and at high titres, confers some virus protection but also reduces host lifespan (Chrostek et al., 2013; Hedges et al., 2008).

As temperatures ideally suited to *D. melanogaster* are generally higher than those suited to *Wolbachia*, we predict that manipulating their host’s behaviour to seek cooler temperatures would be beneficial. Thus, our objective here is to use an established behavioural assay to test the capacity for *Wolbachia* infection to alter host *T*_p_.

## Materials and Methods

*Drosophila melanogaster* from the Oregon RC line were infected with the *w*MelCS line of *Wolbachia pipientis* (hereafter +*Wol*) (Hedges et al., 2008). All flies were reared in 25°C incubators on a standard cornmeal diet and 12 h light/dark cycle. The paired *Wolbachia*-free fly line (hereafter control) was generated from the same *w*MelCS-infected fly line by treating flies with 0.03% tetracycline. We confirmed that the *Wolbachia* strain was *w*MelCS by PCR using two primer sets as described in Riegler et al. (2012). These flies were reared on a standard cornmeal diet for at least five generations before use to recover after tetracycline treatment, and microbiota was reconstituted and standardised following standard procedures. Here, temperature preference assays were conducted on adult male flies aged 4–7 days, but both male and female flies of various ages have been shown to exhibit very similar temperature preferences even when infected by *Wolbachia* (Truitt et al., 2018).

Temperature preference assays used an identical thermal gradient apparatus to that previously described in (Arnold et al., 2015). The apparatus achieved a stable linear gradient of 0.2°C per cm across a temperature range of 17.5–33.5°C (Arnold et al., 2015). Temperatures were measured throughout the experiment by five K-type thermocouples suspended in the gradient airspace, held by bungs that were fitted into an acrylic cover, recorded by a Squirrel 2040 temperature meter.

For each trial, five flies were gently tipped into the centre of the apparatus (25°C) and allowed to freely move about the apparatus for 30 minutes. At the end of the trial period, flies were anaesthetised by CO_2_ that was introduced into both ends of the gradient at a low-flow rate to prevent changes to the position of flies. Distance along the gradient was then used to determine the preferred temperature at the position of rest for each fly in the gradient, *T*_p_. A total of 58 flies for each treatment were assessed for *T*_p_, across 24 replicate trials.

As circadian rhythm affects *T*_p_ in *Drosophila*, we always conducted temperature preference assays between 09:30 and 13:30, a time period across which *T*_p_ is stable (Kaneko et al., 2012). Assays were also conducted in darkness by covering the apparatus in black material to prevent phototactic behaviour affecting positioning.

Exploratory data analyses where flies in each trial were treated as non-independent in a mixed-effects model (trial as a random variable) found the variance component of trial to be essentially zero. We therefore assumed that *T*_p_ of individual flies were independent and not affected by social interactions during the trial itself (see supplementary material of Arnold et al., 2015). Data were not normally distributed, therefore we conducted a non-parametric Mann-Whitney *U* test to compare the preferred temperatures of control and +*Wol* flies. We then calculated Cohen’s *d* with 95% confidence intervals (CIs) as a measure of effect size, given that *p* is not always robust. All analyses were conducted in the R environment for statistical computing (v3.4.1).

## Results and discussion

We found that flies infected with *Wolbachia* strain *w*MelCS preferred cooler temperatures compared to those without any *Wolbachia* (Fig. 1). The difference between populations was significant even at α = 0.01 (Mann-Whitney *U* test; *W* = 2179, *p* = 0.006), which was supported by an effect size and 95% CIs that did not overlap with zero (Cohen’s *d* = 0.394 [0.189 – 0.563]). Both populations exhibited large variance in *T*_p_, ranging between 18.5 and 29°C (Fig. 1A). There is some overlap in *T*_p_ distributions between the populations (Fig. 1B). In absolute terms, +*Wol* flies preferred a cooler mean (± SE) temperature of 25.06 ± 0.25°C compared to the control flies, which preferred 25.78 ± 0.24°C.

**Figure 1.**
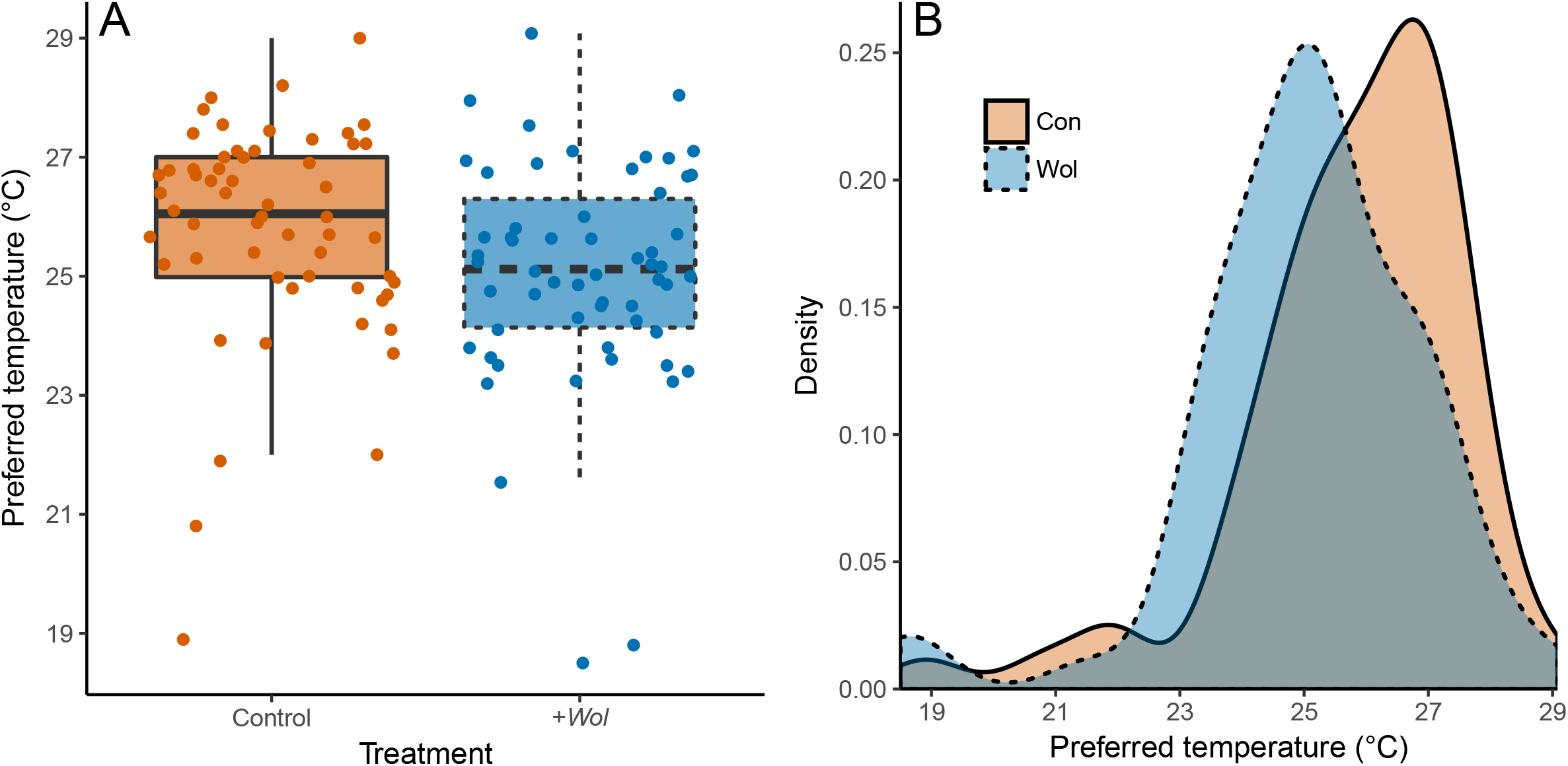
Preferred temperature of uninfected control (orange fill, solid line) and +*Wol* (blue fill, dashed line) *Drosophila melanogaster*. (A) Boxplot of preferred temperature showing median (thick black line), upper and lower quartiles (box), 95% confidence intervals (error bars), and raw data points for control and +*Wol* flies. Each population had *n* = 58 individuals. (B) Density plot showing smoothed distributions of the preferred temperature data for each population.

Our finding is one of the first empirical accounts of *Wolbachia*-infected flies exhibiting a preference for cooler temperatures. Recently, Truitt et al. (2018) demonstrated that three strains of *Wolbachia* (*w*Mel, *w*MelCS, and *w*MelPop) all significantly reduce the *T*_p_ of *D. melanogaster* (strain *w^1118^*). The authors found that *w^1118^* flies infected with *w*MelCS had a 4°C lower mean *T*_p_ compared to uninfected flies, although the *T*_p_ of uninfected *w^1118^* flies was 24.4°C, which is 1.4°C lower than the Oregon RC flies from the present study. We have identified a similar *T*_p_ shift of *Wolbachia*-infected flies in the same direction (although lower magnitude) in a different host strain, using a different temperature preference apparatus. Our results not only support the findings of Truitt et al. (2018), but also suggest that this biologically-interesting phenomenon could be reasonably common.

One explanation for our findings is that *Wolbachia* could potentially manipulate an important aspect of host thermoregulatory behaviour. Pathogens can improve their transmission probability and reproductive capacity by inducing host behavioural changes (Lefèvre & Thomas, 2008), and *Wolbachia* can affect host mate choice and activity levels (van Houte et al., 2013). Changes in thermoregulation behaviour has been well studied from the perspective of the infected host, particularly behavioural fever, where the host elevates its *T*_b_ by behavioural means to rid itself of the pathogen (Kluger, 1979). However, it is less clear whether pathogens manipulate host *T*_p_, especially when the pathogen is not parasitic (i.e., commensalistic or mutualistic) and for decreases to *T*_p_.

Variance of temperatures in nature may lead to populations with mixed or incomplete *Wolbachia* infection (van Opijnen & Breeuwer, 1999), but it is possible that *Wolbachia* could manipulate host *T*_p_ to maximise its own fitness without negatively affecting the host. Cyclic heat stress fluctuating between 26°C and 37°C at 12 h intervals significantly reduced *Wolbachia* density and cytoplasmic incompatibility of *w*Mel and *w*MelPop-CLA, but not *w*AlbB in *Aedes aegypti* (Ross et al., 2017). The *w*MelCS strain used in the present study likely shares a recent field origin with *w*Mel (Riegler et al., 2005), which is a widely used dengue-suppressing *Wolbachia* strain that itself also alters host *T*_p_ to prefer cooler temperatures (Truitt et al., 2018). Temperature fluctuations like the cyclic heat stress experiment by Ross et al. (2017) might well be experienced naturally in tropical regions. This would likely result in incomplete infection and reduced transmission success, which could explain the erratic temporal and spatial dynamics of *Wolbachia* spread in controlled infected-vector release programs (e.g., Schmidt et al., 2017). Our finding suggests that *Wolbachia*-infected hosts prefer cooler temperatures and might be likely to seek out cooler microclimates, which would reduce exposure to and lessen the fitness consequences of high temperatures.

An alternative hypothesis is that flies infected with *Wolbachia* exhibit a behavioural chill to restrict pathogen replication. Infection with *Pseudomonas aeruginosa* lowered the *T*_p_ of *D. melanogaster*, but this behavioural chill response was inefficient and did not reach the critical temperature that would increase survival and limit bacteria growth rate (Fedorka et al., 2016). The change *T*_p_ that we observed in flies infected with *w*MelCS could also be a host behavioural response. However, as discussed earlier, *Wolbachia* tends to perform worse at higher temperatures than are optimal for *D. melanogaster*, and therefore we suggest that manipulation of the host *T*_p_ by the pathogen is more likely.

The absolute decrease in *T*_p_ of less than 1°C that we observed in flies infected with *Wolbachia w*MelCS provides little buffer to the predicted 2–4°C increase by 2100 due to climate change. However, *Wolbachia* are maternally inherited and exposure to high temperatures can reduce vertical transmission in only a few generations (Corbin et al., 2016). If the infected host prefers cooler temperatures, then this behaviour would confer a selective advantage for *Wolbachia*. Arthropod hosts of *Wolbachia* have rapid generation times relative to the forecast rate of temperature increase, therefore it is conceivable that a minor change in *T*_p_ could be enhanced by selection across generations to allow continued transmission and mitigate fitness consequences.

This study paves the way for discovering the mechanisms by which *Wolbachia* - infection alters host *T*_p_. Whether the observed phenomenon is due to *Wolbachia* directly manipulating host behaviour, a host defense response, or a by-product of infection will need to be determined. The efficacy of introductions of populations of *Wolbachia*-infected vectors may hinge upon a better understanding of complex host-pathogen-environment interactions. Testing for *Wolbachia*-induced changes in thermal preference across multiple host and pathogen strains will elucidate whether unexpected ecological and evolutionary responses might occur in planned vector releases in a changing climate.

## Acknowledgements

We thank Prof. Craig R. White for providing laboratory space and equipment for this study, and constructive comments from three anonymous reviewers that improved this manuscript.

## Contribution of authors

PAA established the methodology, lead the analyses, interpreted results, and wrote the manuscript; SL and ALS collected the data and contributed to the manuscript draft; KNJ supervised the project, interpreted results, and contributed substantially to the manuscript draft.

